# The role of mu-opioid receptors in pancreatic islet alpha cells

**DOI:** 10.1101/2024.05.13.593899

**Authors:** Chen Kong, Daniel C. Castro, Jeongmin Lee, David W. Piston

**Affiliations:** Department of Cell Biology & Physiology, Washington University School of Medicine, St Louis, Missouri, 63110, USA; Mallinckrodt Institute of Radiology, Washington University School of Medicine, St Louis, Missouri, 63110, USA

**Author notes:** C. K. and D.C.C. contributed equally to this work. Corresponding authors: Daniel C. Castro,; David W. Piston.

**Keywords:** Human islets, OPRM1, MOPR, Glucagon secretion, Alpha-cell, Beta-cell

## Abstract

30% of people in the United States have diabetes or pre-diabetes. Many of these individuals will develop diabetic neuropathy as a comorbidity, which is often treated with exogenous opioids like morphine, oxycodone, or tramadol. Although these opioids are effective analgesics, growing evidence indicates that they may directly impact the endocrine pancreas function in human and preclinical models. One common feature of these exogenous opioid ligands is their preference for the mu opioid receptor (MOPR), so we aimed to determine if endogenous MOPRs directly regulate pancreatic islet metabolism and hormone secretion. We show that pharmacological antagonism of MOPRs enhances glucagon secretion, but not insulin secretion, from human islets under high glucose conditions. This increased secretion is accompanied by increased cAMP signaling. mRNA expression of MOPRs is enriched in human islet α-cells, but downregulated in T2D islet donors, suggesting a link between metabolism and MOPR expression. Conditional genetic knockout of MOPRs in murine α-cells increases glucagon secretion in high glucose conditions without increasing glucagon content. Consistent with downregulation of MOPRs during metabolic disease, conditional MOPR knockout mice treated with a high fat diet show impaired glucose tolerance, increased glucagon secretion, increased insulin content, and increased islet size. Finally, we show that MOPR-mediated changes in glucagon secretion are driven, in part, by K_ATP_ channel activity. Together, these results demonstrate a direct mechanism of action for endogenous opioid regulation of endocrine pancreas.

## INTRODUCTION

Diabetes and prediabetes affects tens of millions of people in the United States [1]. While effective therapies exist for some patient populations, pathologies related to long-term dysregulation of glycemia remains a major medical problem. Further complicating the diabetes prognosis are emergent comorbidities such as diabetic neuropathy, which has historically been treated with analgesic opioids like morphine, oxycodone, or fentanyl [2]. Opioids are well known for their analgesic and recreational properties, but an under-discussed feature is that opioids appear to directly influence glucose homeostasis [3–6]. Although metabolic disturbance is not typically associated with opioid use, the intersection of the opioid, diabetes, and pain epidemics poses a real, albeit complicated, threat to human health.

Endogenous and exogenous opioids have been shown to dysregulate insulin secretion when injected systemically or directly applied to pancreatic islets [5], primarily via their actions on the inhibitory Gi-coupled mu opioid receptor (MOPR) [7]. When stimulated, MOPRs induce hyperglycemia although specific drugs may differ in their effects depending on the satiety of the organism [8–10]. Even though opioid stimulation *in vivo* increases circulating glucose, it acutely increases both insulin and glucagon secretion in *ex vivo* isolated islets. Consistent with the varied effects that opioids have on glucose and hormones, disruption of MOPR (e.g., via naloxone) dysregulates metabolism in ways that are both similar and opposite to opioid stimulation [11–13]. One example linking MOPRs and pancreatic islet function [14] showed that MOPR constitutive knockout mice displayed increased body weight, enhanced glucose tolerance, and a 4-fold increase in insulin release. They showed that MOPRs are expressed in pancreatic islets, and that isolated islets from MOPR knockout mice were unusually sensitive to glucose-stimulated insulin secretion. Despite several studies showing increased glucagon in combination with enhanced insulin secretion [15], limited literature exists regarding the effect of the opioid system on islet α-cells or glucagon secretion. This suggested to us that opioids might directly affect glucagon secretion, and thereby overrule the suppressing effects of insulin on glucagon secretion.

To test this hypothesis, we sought to determine how MOPRs regulate glucagon secretion from α-cells in human and mouse primary islets. We report that pharmacological antagonism of MOPRs increases glucagon secretion, but not insulin secretion, from human primary islets. This effect is most pronounced in high glucose conditions. mRNA expression and *in situ* expression experiments further corroborated the presence of endogenous MOPRs on primary islets. Notably, MOPR expression is reduced in humans with Type 2 diabetes or obese mice, suggesting an interaction between metabolic state and opioid activity. Specific genetic deletion of *Oprm1* (opioid receptor mu 1) from the α-cells results in impaired glucose tolerance and augmented insulin production after high fat diet. Finally, we show that the secretory effects of MOPR antagonism are driven by the recruitment of K_ATP_ channels, identifying a potential mechanism of action. These data identify a novel opioid system that has major implications for many disease states, including diabetes, obesity, opioid use disorder, and pain.

## MATERIALS AND METHODS

### Animals and plasma islet hormones

All animal experiments were performed under approval of the Washington University Institutional Animal Care and Use Committee (protocol # 20-0381). GCGCre *Oprm1*^fl/fl^ mouse line was generated by multiple crosses of OPRM1 loxP mice (obtained from Jackson Laboratory stock No: 030074) and mice expressing Glucagon-iCre (Jackson Laboratory stock No:030663). GCG-Cre+ *Oprm1*^fl/fl^ mice lack OPRM1 specifically on α-cells (KO) and their littermates GCG-Cre-*Oprm1*^fl/fl^ mice (Control) were used for the experiments. Mice were genotyped for iCre and OPRM1 genes via ear punch tissue by Transnetyx (Cordova, TN USA). Mice were fed a standard rodent diet (13.2% calories from fat) or a moderately high fat diet (21.6% calories from fat). Mice were used at 10 to 18 weeks of age or 12 weeks after the high fat diet feeding for the experiments. The body weight and blood glucose were monitored weekly. Plasma levels of insulin and glucagon were measured by ELISA (Crystal Chem, #90080 and #81520 respectively).

### Glucose and Insulin Tolerance Test

Mice were fasted for 6 h followed by an intraperitoneal injection of 2 g/kg body weight of sterile D-glucose solution for glucose tolerance test (GTT) or fasted for 4 h before receiving 0.5 unit/kg body weight of human recombinant insulin (Novolin 70/30, U-100) for insulin tolerance test (ITT). Whole blood was obtained from the tail vein at the indicated time points (0, 15, 30, 60, 90, and 120 min) after glucose injection. Blood glucose levels were measured using a Glucocard Vital blood glucose monitor (Arkray). To measure the plasma glucagon and insulin levels the mice were fasted for 6 h followed by an intraperitoneal injection of glucose (2g/kg body weight) in separate time. Blood was collected in EDTA-containing tubes, centrifuged (2500g, 15 min at 4°C) to collect the plasma.

### Preparation of isolated islets and dispersed cells

The mouse pancreas was surgically removed, then digested in 9ml of HBSS (Gibco, #14025134) with 2 mg/ml collagenase P (Roche, #11213873001) in the tube rotating at 40 rpm for 35 minutes at room temperature and washed two times in HBSS with 1% BSA. The cell suspension was filtered by a cell strainer. Islets were manually picked under a stereomicroscope and allowed to recover overnight in RPMI 1640 (Gibco, #11879020) supplemented with 10% FBS, 11 mM glucose, and 100 U/ml penicillin-streptomycin. For the islet cell dispersion, the isolated Islets were washed HBSS and then dissociated in Accutase (Innovative Cell Technologies, # AT-104) for 15 min at 37°C with intermittent trituration. Dissociated cells were resuspended in islet media for subsequent use.

### Langerhans islet planar area measurement and histology

Islets from α-cell OPRM1 KO and control mice were isolated as described above. A picture for each group of islets isolated from a single animal was photographed using a Nikon TS100 microscope. The planar area of islets with normal morphology was measured using the ImageJ software (National Institutes of Health, Bethesda, MD).

### Secretion assays

The isolated islets were equilibrated in RPMI1640 with 10 mM HEPES, 0.1% BSA and 2.8 mM glucose in groups of 6 to 8 size matched islets per tube (human islets in groups of 5-6 per tube). The supernatant was collected after 1 h incubation at 37 °C (equilibration), then treated in 1 or 11 mM glucose with or without treatments (DAMGO 1µM, CTAP 1µM or Tolbutamide 300 µM) for 1 h at 37 °C and the supernatant was collected (secretion). For dispersed islet secretion, dispersed islet cells were placed in microcentrifuge tubes (10^4^ cells/assay) and equilibrated in 5.5 mM glucose for 1 h at 37 °C. The cells were spun down and resuspended in different treatments and supernatant was collected. The data represent the secretion concentration of glucagon or insulin in each tube normalized by the equilibration factor (equilibration concentration divided average equilibration concentration in the islet samples used for the experiment from the same mouse). Insulin and glucagon concentrations were measured by Lumit Immunoassay kit (Promega # W8022).

### Islet hormone content

The total islet hormone content from each secretion assay was extracted overnight with an acid-ethanol (1.5% 12 N HCl in 70% ethanol) solution in -20°C and measured by Lumit Immunoassay kit. The total islet protein content for each secretion assay was measured with Micro BCA™ Protein Assay Kit (ThermoFisher, #23235). Data represent the glucagon or insulin content per µg of protein in each tube.

### Immunohistochemistry

The mouse pancreases were fixed in 10% buffered formalin solution overnight at room temperature, and the paraffin-embedded pancreatic were sectioned at the Anatomic and Molecular Pathology Core labs in Washington University in St Louis. For immunohistochemistry assays, the pancreas sections were first deparaffinized with sequential incubation in xylene (10minutes, twice), 100% ethanol (5 minutes), 95% ethanol (5 minutes), 70% ethanol (5 minutes) and deionized water (2 minute). Antigen retrieval was conducted by boiling the pancreas slices in Diva Decloaker buffer (Biocare, #DV2004LX) in a hot water bath at 98 °C for 20 minutes, followed by natural cooling down to below 40°C, and washing with deionized water. The tissue slices were permeablized with 0.1% Triton X-100 in PBST (PBS containing 0.05% Tween20) for 20 min at room temperature. After 1h incubation in blocking buffer (1% BSA and 5% goat serum in PBST), slices were incubated overnight at 4 °C with primary antibodies against glucagon (AvantGen, #DA-2013) followed by incubation with the corresponding secondary antibody that were conjugated to various fluorescent agents. For nuclear staining, DAPI (Sigma, # MBD0015) was added to the slides after secondary antibody incubation and incubated for 10 minutes at room temperature. Slices were then washed in PBS and mounted with ProLong Glass Antifade Mountant (Invitrogen, #P36980) and processed for fluorescence microscopy (Zeiss LSM880).

### Fluorescence *in situ* hybridization

RNAScope Fluorescence Multiplex Version 2 Assay combine with immunofluorescence on formalin-fixed paraffin-embedded (FFPE) tissue sections was performed according to the manufacturer’s instructions (ACDBio, Cat. No. 323100). Briefly, mouse pancreas FFPE tissue sections were deparaffinized, antigen retrieved, hybridized with positive (ACD, 320881), negative (ACD, # 320871) and Mm-Oprm1-O4-C2 (ACD, #544731) probes and developed with Opal-520 (FP1487001KT). Slices were then immunostained with glucagon antibody (Avantgen, DA-2013) as an α-cell marker. The pancreas sections were imaged on the Zeiss LSM880 fluorescence microscope (Nikon, Ti-E).

### Immunofluorescence

Dispersed islet cells were cultured overnight on glass bottom chambers coated with 2µg/cm^2^ rhLaminin-521(Gibco, # A29248). The cells were fixed at room temperature with 4% paraformaldehyde for 20 min followed by permeabilization with 0.1% Triton X-100 in PBST (PBS containing 0.05% Tween 20) for 5 min at room temperature. After 1h incubation in blocking buffer (1% BSA and 5% goat serum in PBST), the cells were incubated overnight at 4 °C with primary antibodies against glucagon (AvantGen, #DA-2013), Cre (Cell Signaling #15036) in PBST with 1%BSA followed by 1h incubation with secondary antibodies and DAPI staining for nuclear counterstain. Cells were imaged on the Zeiss LSM880 fluorescence microscope (Nikon, Ti-E).

### Human islet culture

Human islets were cultured in CMRL medium (Gibco 11530-037) supplemented with 10% FBS, 2 mM glutamine, 100 U/mL penicillin and100 mg/mL streptomycin at least one day before the experiments (qPCR, intact islet or dispersed islet cell secretion and immunofluorescence). Human pancreatic islets were provided by the NIDDK-funded Integrated Islet Distribution Program (IIDP) (RRID:SCR_014387), NIH Grant #UC4DK098085, donor information provided in Table S1.

### Islet cAMP FRET

Pancreatic islets were cultured on 8 well-chambered cover glass systems (Cellvis, C8-1,5H-N) coated with Rh-laminin521 for 2 days. Islets were incubated with adenovirus expressing pEpac-SH187 [16] a cAMP sensor in living cells for 3 days prior to experiments. For the FRET imaging, seeded islets were placed in Zeiss LSM 880 confocal microscope with heated, CO_2_-controlled microscope sage (37C, 5% CO_2_) with Plan-Apochromat 20X/ 0.2 M27 objective. Islets were equilibrated for 30 minutes with KRBH with 0.1% BSA with 2 mM glucose. Islets were incubated with KRBH with 0.1% BSA with 1mM or 11mM glucose for 30 minutes and a baseline image was acquired. Then, the buffer was switched to KRBH with 0.1% BSA and either 1mM or 11mM glucose with treatment and incubated for 30 minutes prior to acquiring a treatment image. For identification of α-cells, islets were treated with epinephrine (10µM) in11mM glucose for 10 minutes, and each cell’s FRET ratio was calculated to distinguish between α and β-cells; α-cells showed increased cAMP, while β-cells exhibited the opposite response [17]. Signals for acceptor and donor (mTurquoise, mVenus) were measured with LSM 880 spectral detector mode with 405nm excitation and detected at an emission range of 445-650nm. The acceptor and donor signals were acquired through linear unmixing with the ZEISS ZEN imaging software from the entire emission spectrum image. Since the FRET construct is inversely proportional to cAMP-EPAC level, the Donor-to-acceptor ratio was calculated, and before and after treatment, the percentage change was calculated.

### Quantitative RT-PCR analysis

Total RNA from mouse and human islets was extracted using a RNeasy micro kit (Qiagen # 74004) and reverse transcribed using a SuperScript™ VILO™ Master Mix (Invitrogen # 11755050). Quantitative PCR was performed with Power SYBR Green PCR Master Mix (Applied Biosystems # A25742) on ViiA7 qPCR machine (Applied Biosystems). All the results were analyzed using the delta-delta-Ct method and normalized to18s and representative of three biological replicates and two technical replicates. Primer sequences used are Human OPRM1: 5’ GCCTCAACCCAGTCCTTTAT; 3’ GTTGGAAGAGGTTGGGATACA. Mouse OPRM1: 5’ GCCCTCTATTCTATCGTGTGTG; 3’ GTAGATGTTGGTGGCAGTCTT. iCre:5’ GGCTGTGAAGACCATCCAACA; 3’ CACACCAGACACAGAGATCCA. 18s: 5’-GTAACCCGTTGAACCCCATT; 3’-CCATCCAATCGGTAGTAGCG

### Data Analysis and Statistics

Data were analyzed with Microsoft Excel and ImageJ. All statistical analyses were performed in Prism 9 software (GraphPad Software, La Jolla, CA) using unpaired two-tailed Student’s t-test. Data are presented as pooled data (n refers to numbers or sample numbers as indicated in figures) as mean ± SEM, and *P*<0.05 was considered significant.

## RESULTS

### OPRM1 expression and opioid effects on glucagon secretion from human islets

To establish that *Oprm1* (Opioid Receptor Mu 1) is expressed in both human and murine islets, we used qPCR to quantify the relative expression of the *Oprm1* gene (Figure 1A). *Oprm1* is more highly expressed in human than mouse islets. In human islets, *Oprm1* expression is comparatively lower in Type-2 Diabetes. Similarly, relative expression of *Oprm1* in obese leptin receptor deficient mice (LepR db/db) is downregulated. These results suggest that MOPRs in endocrine pancreas are dynamically regulated in disease states.

**Figure 1.**
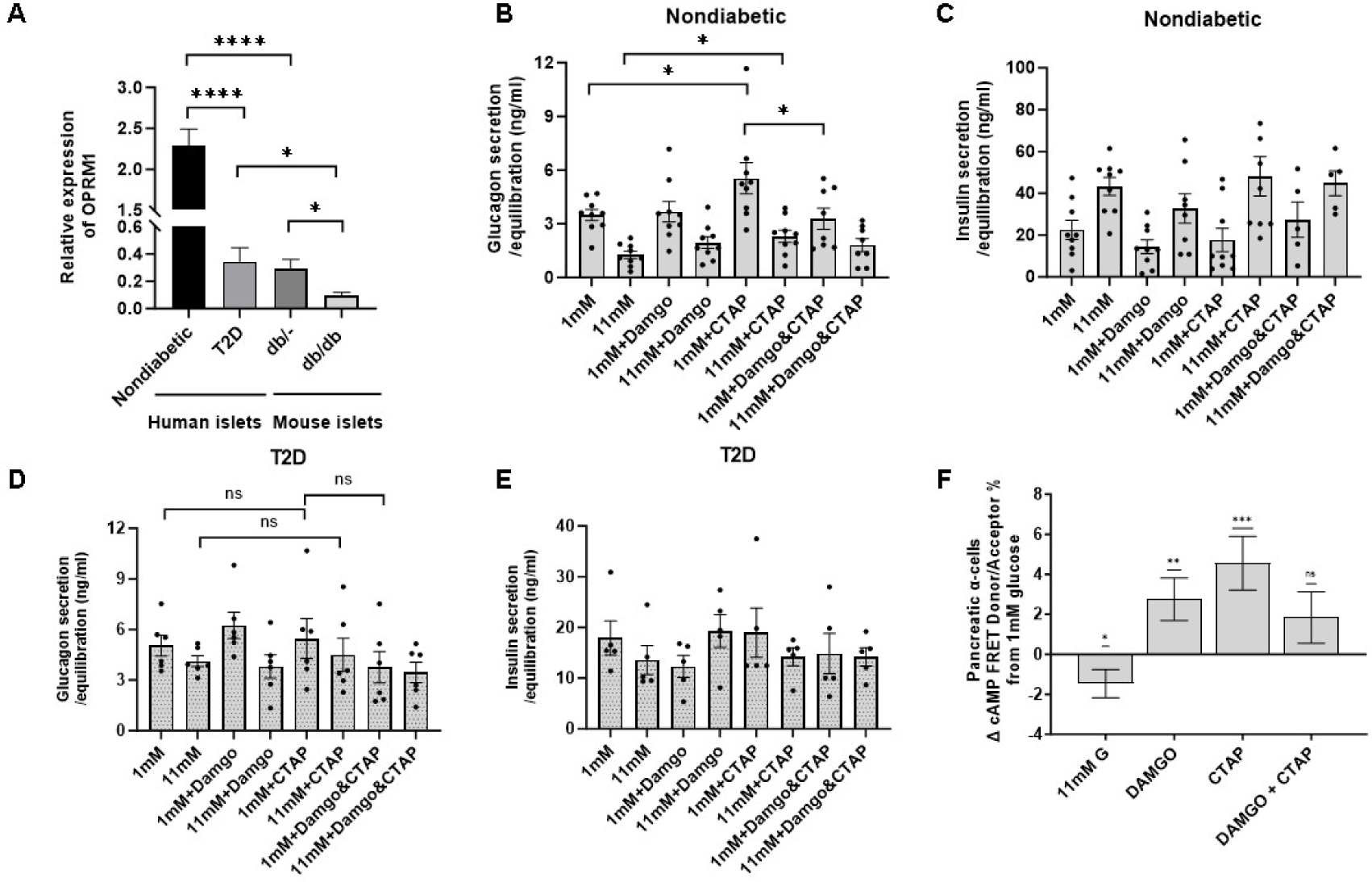
OPRM1 expression and islet secretion in in human islets. (A) OPRM1mRNA expression in islets from nondiabetic (n=4) or type 2 diabetes (T2D, n=3) human islets and Lepr db/-BKS (control n=4) or Lepr db/db BKS (T2D model, n=4) mouse islets with qPCR. Data are presented as OPRM1 expressed relative to the housekeeping gene 18S. Human islet glucagon (B) and insulin (C) secretion treated by receptor agonist DAMGO and antagonist CTAP: CTAP treatment increased glucagon but not insulin secretion (n = 9 experiments, using islets from 6 donors). Human T2D islet glucagon (D) and insulin secretion (E) with treatment of DAMGO and CTAP: No significant change has been shown with CTAP treatment (n = 6 experiments, using islets from 3 donors). (F): Epac-cAMP FRET at low (1mM) and high (11mM) glucose and treatment of DAMGO, CTAP or both in the islets from non diabetic donors (n = 3 experiments, using islets from 2 donors). cAMP level increased in alpha cells when the islets treated with DAMGO and CTAP. Data represented as mean ± SEM. *, *P* 0.05; **, *P* 0.01; ***, *P* 0.001; ****, *P* 0.0001.

To establish a functional link between MOPR expression and endocrine hormone secretion, human donor islets were treated with the MOPR selective agonist DAMGO or the MOPR selective antagonist CTAP. Glucagon and insulin secretion was measured in 1 mM (low) and 11 mM (high) glucose conditions. As expected, baseline glucagon secretion is higher in low glucose conditions than high glucose (Figure 1B). MOPR stimulation via DAMGO shows no effect on glucagon secretion in either low or high glucose conditions, although there is a non-significant trend towards increased glucagon secretion under high glucose conditions. In contrast, MOPR antagonism via CTAP increases glucagon secretion in both low and high glucose conditions. Co-administration of DAMGO and CTAP blocks the CTAP alone effects. Unlike glucagon secretion, insulin secretion is unaffected by either DAMGO or CTAP, indicating that MOPRs may primarily exert their effects on α-cells (Figure 1C).

We next sought to test whether pathological downregulation of *Oprm1* gene expression impacts MOPR stimulated effects on hormone secretion. We pharmacologically agonized or antagonized MOPRs and measured glucagon or insulin secretion in Type 2 diabetes human donor islets. As expected, suppression of glucagon secretion in high glucose conditions is impaired in T2D donor islets (Figure 1D) [18]. Unlike healthy donor islets (Figure 1B), CTAP antagonism had no effect on glucagon secretion, regardless of low or high glucose conditions (Figure 1D). Similarly, DAMGO stimulation had no effect on glucagon secretion, and neither CTAP nor DAMGO impacted insulin secretion (Figure 1E).

Finally, to test whether changes in MOPR activity would influence α-cell activity, we used a FRET-based cAMP biosensor to measure cAMP levels during live cell imaging before and after MOPR agonism (DAMGO) or antagonism (CTAP) (Figure 1F). Islet cell-types were identified by their response to epinephrine (10µM) in 11mM glucose. As expected, cAMP decreases when the glucose concentration changed from low (1mM) to high (11mM). In contrast, stimulation via DAMGO or blockade via CTAP increases cAMP in low glucose conditions. Co-administration of DAMGO and CTAP does not significantly impact cAMP. In high glucose conditions, neither DAMGO nor CTAP alone significantly modulate cAMP. However, co-administration of the drugs results in increased cAMP. Together, these results suggest that MOPR expression is dynamically modulated by disease state, and that modulation of MOPR in isolated islets can modulate cellular and hormonal activity.

### Generation of an α-cell–specific *OPRM1* KO mouse model

Expression, secretion, and signaling results suggest that MOPRs primarily exert their effects by modulating glucagon secretion from α-cells. To determine the role of MOPRs in α cells specifically, we selectively deleted MOPRs from α-cells by crossing GCG-Cre mice with *Oprm1*^fl/fl^ mice. *Oprm1*^fl/fl^ mice possess *loxP* sites flanking exons 2-3 of the opioid receptor mu1 (*Oprm1*) gene (Jackson Laboratory), and have been used to generate selective knockouts with other cre lines [19,20]. One potential limitation to this approach is that the proglucagon gene can target cell populations outside of the pancreas, such as the distal ileum and the large intestine, as well as certain brain cells. To verify that our knockdown was selective to pancreatic α-cells, we used qPCR to measure the relative Cre mRNA across tissue samples. Cre expression is nearly undetectable in brain, colon, and liver from conditional Knockout control mice. In contrast, we observe a significant amount of Cre expression in islets isolated from GCG-Cre+ *Oprm1*^fl/fl^ mice (Figure 2A). Follow up studies using immunofluorescence to look at protein expression on dispersed islet cells similarly show that Cre expression is localized to the α-cells of the GCG-Cre+ *Oprm1*^fl/fl^ mice (Figure 2B). After confirming the selective expression of Cre, we next tested whether conditional deletion of MOPRs was limited to Cre-expressing α-cells. Here, we quantified *Oprm1* mRNA on pancreatic islets using RNAscope combine with immunofluorescent staining for glucagon. RNAscope control experiment is shown in the Supplemental figure 1. Our results show that while *Oprm1* is expressed on both α- and putative β-cells in Cre-control mice, GCG-Cre+ *Oprm1*^fl/fl^ mice show selective deletion in only the α-cells (Figure 2C). This data is quantified by measuring the number of RNAscope labeled puncta on each cell (Figure 2D). In sum, our results indicate that we have developed a mouse line that selectively disrupts *Oprm1* expression in pancreatic α-cells.

**Figure 2:**
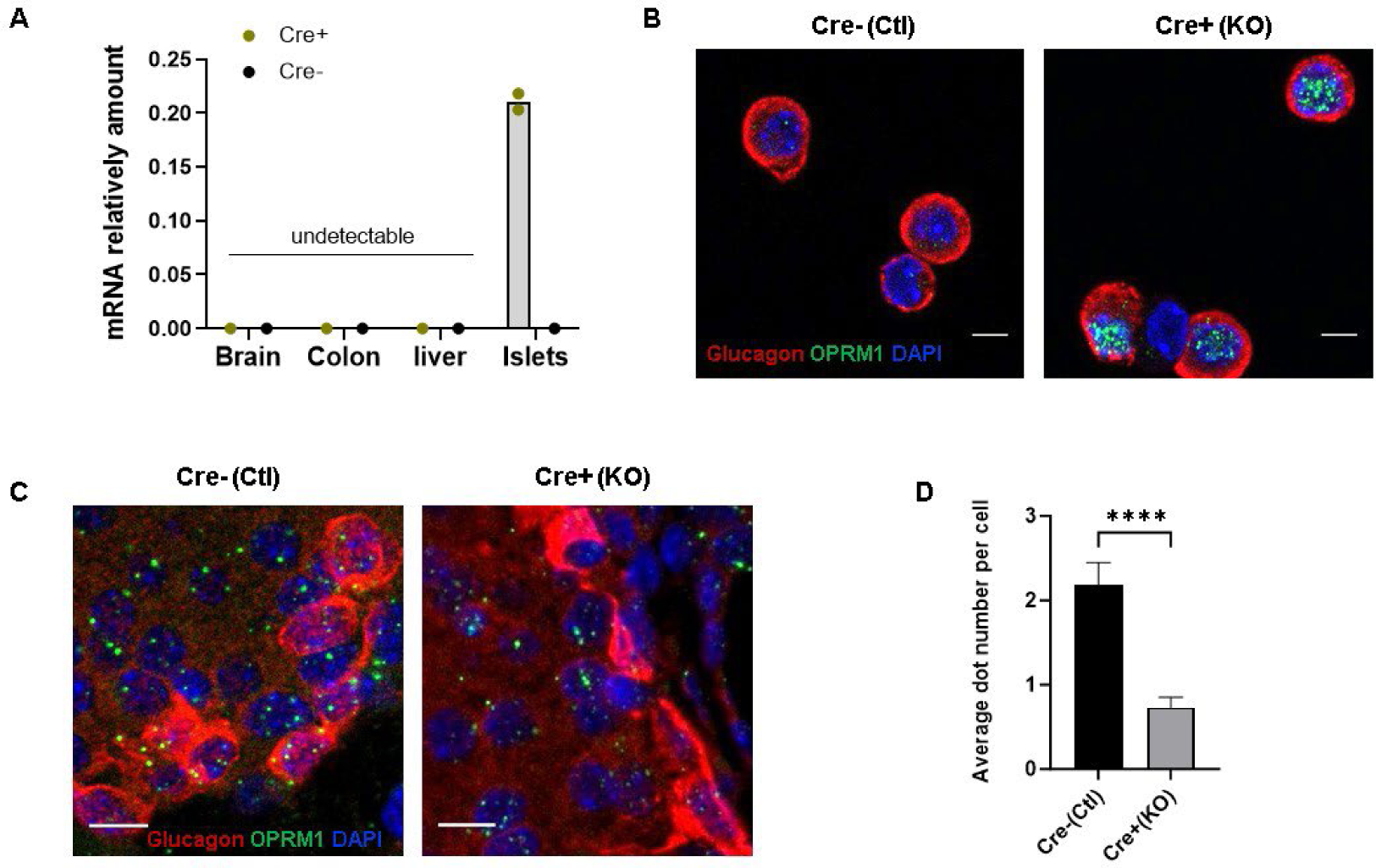
Verification of islet alpha cells-specific deletion of OPRM1 gene in GCGCrefl/fl mouse mode. (A) iCre gene expression relative to 18s (housekeeping control) with qPCR was only detected in islets from αKO mice, undetectable in brain, colon and liver in both control and KO mice and islets in the control mice. (n = 3 mice per genotype.). (B) Cre expression in islet alpha cells. Dispersed islet cells from Cre-(control) and Cre+ (KO) mice were immunostained with Cre and glucagon antibodies. Cre staining (Green) was merged with glucagon staining (Red), an alpha cells marker and DAPI nuclear staining (Blue). Scale bars, 5 μm. (C) RNAscope ISH for OPRM1 RNA expression in pancreatic alpha cells was performed by using OPRM1 probe (green) with immunostaining of glucagon (red) co-detection and nuclei were counterstained with DAPI (blue) in pancreas sections from KO and control mice. Scale bars, 5 μm Images are representative of three experiments with three mice each. (D) Quantification of RNAscope® assay shows the average dot number per cell using the Image J software. N > 15. Data represented as means ± SEM. *, *P* < 0.05; **, *P* < 0.01; ***, *P* < 0.001; ****, *P* < 0.0001.

### OPRM1 deletion from α-cells disinhibits glucagon secretion under high glucose conditions

We assessed the metabolic phenotype of *Oprm1* deletion from α-cells by monitoring KO mice and littermate controls for body weight, insulin tolerance, plasma glucose, glucagon, and insulin levels. We did not observe obvious phenotypic differences between Cre^+^ and Cre^-^ mice (Supplementary Figure 2). However, isolated KO islets show increased glucagon secretion under high glucose conditions compared to control islets (Figure 3A). Hormone content for glucagon (Figure 3B) or insulin secretion (Figure 3C) do not show any difference across glucose concentration conditions between Cre^+^ and Cre^-^ mice. These results indicate that loss of MOPRs on α-cells results in disinhibition of glucagon cells, rather than overproduction of glucagon peptide. Chronic disinhibition of α-cells has previously been shown to impact the progression of a variety of metabolic disorders, suggesting that these mice may be more sensitive to disease states like obesity or T2D. Finally, we note that neither MOPR agonism nor antagonism have a robust effect on glucagon or insulin secretion (Supplementary Figure 3). This could, in part, be due to the comparatively low expression of MOPRs on mouse versus human islets.

**Figure 3.**
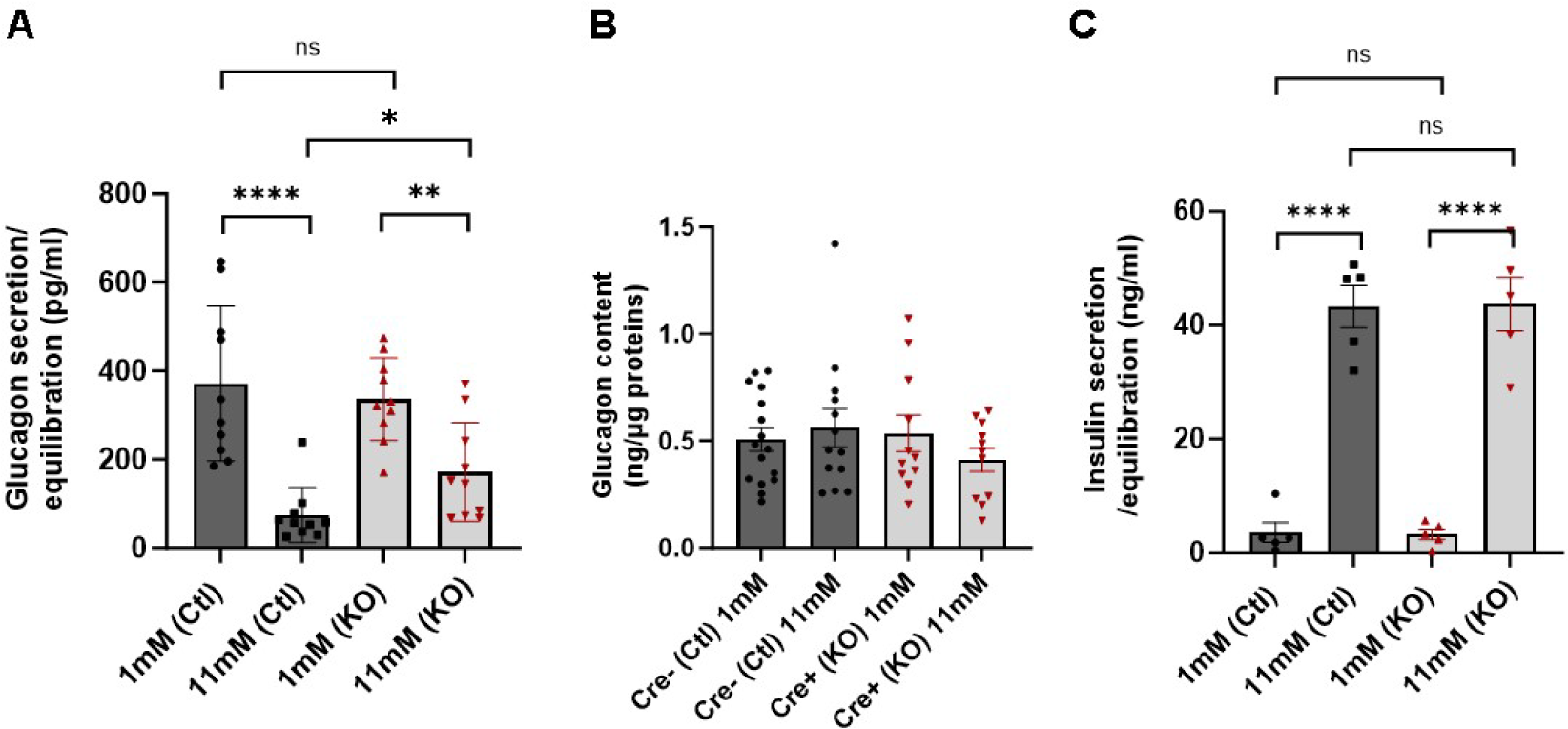
Islet phenotypes in chow diet mice. Isolated Islet secretion from α cell OPRM1 deficiency (KO) and control (Ctl) mice were treated with low glucose (1mM) and high glucose (11mM). (A) Glucagon secretion by KO islets shows similar in the low concentration glucose, less glucose suppression in the high concentration glucose compared to the secretion in control islets and (B) insulin secretion was not altered. (C) KO islets have normal glucagon content normalized with total protein amount in the same batch of islets. Data represented as means ± SEM. *, *P* < 0.05; **, *P* < 0.01; ***, *P* < 0.001; ****, *P* < 0.0001.

### α-cell specific *OPRM1* KO mice develop glucose intolerance and prediabetes phenotypes with high fat diet

Our results from chow fed mice suggest that *Oprm1* deletion in α-cells may predispose mice to hyperglucagonemia or other similar metabolic disturbances. To test this hypothesis, we tested mice on a series of *in vivo* and *in vitro* assays following high fat diet feeding (HFD, 21.6% calories from fat). After 12 weeks of HFD, there were no differences in body weight between control and KO mice (Supplementary Figure 4). However, consistent with our hypothesis that α-cell mediated metabolic activity is augmented by HFD, we observe that KO mice exhibit impaired glucose tolerance compared to Cre^-^ control mice (Figure 4A). Similarly, Cre^+^ mice have higher plasma glucagon levels 35 minutes after glucose administration compared to Cre^-^ control mice (Figure 4B). These results do not appear to be driven by changes in plasma insulin nor by any impairment in the insulin tolerance test, both of which remain normal for both Cre^+^ and Cre^-^ mice (Figure 4C and D). To determine whether the *in vivo* phenotypes could be explained by complementary changes in pancreatic islets, we assayed isolated islets for glucagon and insulin secretion. As expected, islets from Cre^-^ control mice show glucagon secretion similar to wild-type islets (Figure 4E). In contrast, Cre^+^ islets show impaired glucagon secretion at low glucose levels, and enhanced glucagon secretion at high glucose levels (Figure 4E). While secretion patterns differ between Cre^-^ and Cre^+^ islets, total glucagon content is unchanged (Figure 4F). This suggests that high-fat diet dysregulates α-cell activity to release chronically stable levels of glucagon in cKO mice, rather than by disrupting glucagon availability during high or low glucose states. Surprisingly, while glucose-stimulated insulin secretion is similar between islets from Cre^-^ and Cre^+^ mice (Figure 4G), Cre^+^ islets show greater insulin content (Figure 4H) and increased islet size (Figure 4I). These data indicate that specific genetic deletion of *Oprm1* from α-cells predisposes mice to develop a prediabetic phenotype under a HFD challenge. This is consistent across the *in vivo* changes in glucagon secretion and glucose tolerance, as well as the *ex vivo* dysregulation of glucagon secretion and insulin peptide content.

**Figure 4.**
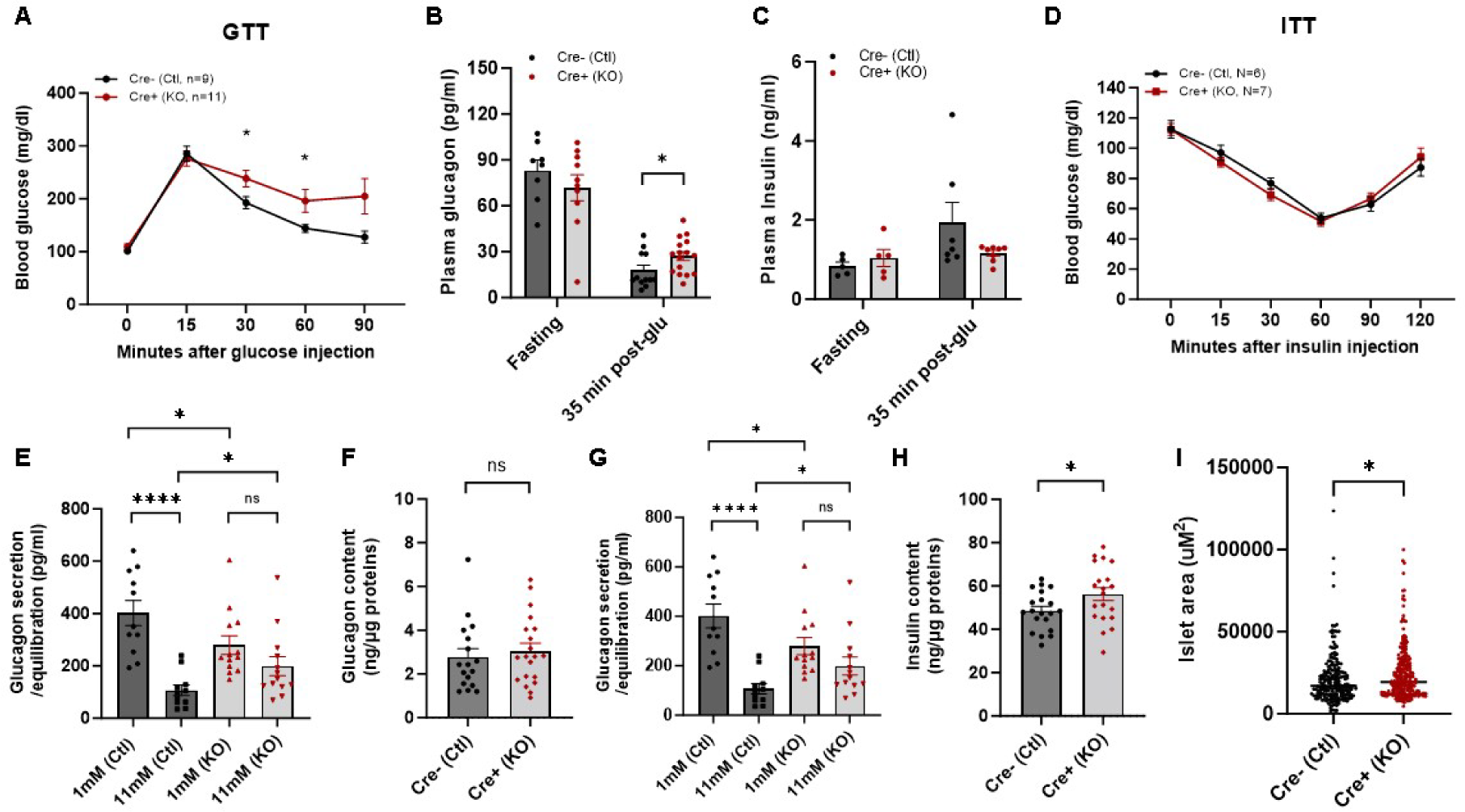
α-cell OPRM1 deletion mice (KO) develop abnormal glucose tolerance and glucagon secretion induced by feeding a high-fat diet. α-cell OPRM1 KO and control mice after 12 weeks on a moderately high fat diet (21.6% calories from fat). (A) Impaired glucose tolerance (n=9 control and 11 KO) and (B) Higher plasma glucagon level and (C) unchanged insulin level at 35 min after administration of glucose in the KO mice compared with control mice. (D) No effect on insulin sensitivity by insulin tolerance testing. (E) Glucagon secretion in both low (1mM) and high (11mM) glucose condition. There was normal in pancreatic glucagon content (F). Insulin secretion trends high in KO islets but does not reach significance (G), and insulin content (H) and larger islets (I) in KO mice compared with controls. Data represented as means ± SEM. Data represented as means ± SEM. *, *P* < 0.05; **, *P* < 0.01; ***, *P* < 0.001; ****, *P* < 0.0001.

### K_ATP_ channels play a role in MOPR regulation of glucagon secretion

Our data indicate that loss of MOPR signaling increases glucagon secretion, particularly in high glucose conditions when glucagon secretion should normally be low. A major mediator of glucose regulated glucagon secretion are K_ATP_ channels [25]. Since MOPRs have previously been shown to produce hyperpolarization in the central nervous system through K_ATP_ channel activation [21–24], we hypothesized that this might be one potential mechanism through which MOPRs modulate glucagon secretion. To test this hypothesis, we treated islets from Cre^+^ and Cre^-^ mice with the K_ATP_ channel blocker tolbutamide. Consistent with previous literature [25–27], tolbutamide decreases glucagon secretion at 1 mM glucose in Cre-control mice (likely due to a reduction in positive current). In contrast, tolbutamide does not augment or disrupt glucagon secretion in Cre^+^ mice, suggesting that chronic reduction in MOPR leads to a functional downregulation of K_ATP_ channels, regardless of the glucose condition (Figure 5A).

**Figure 5.**
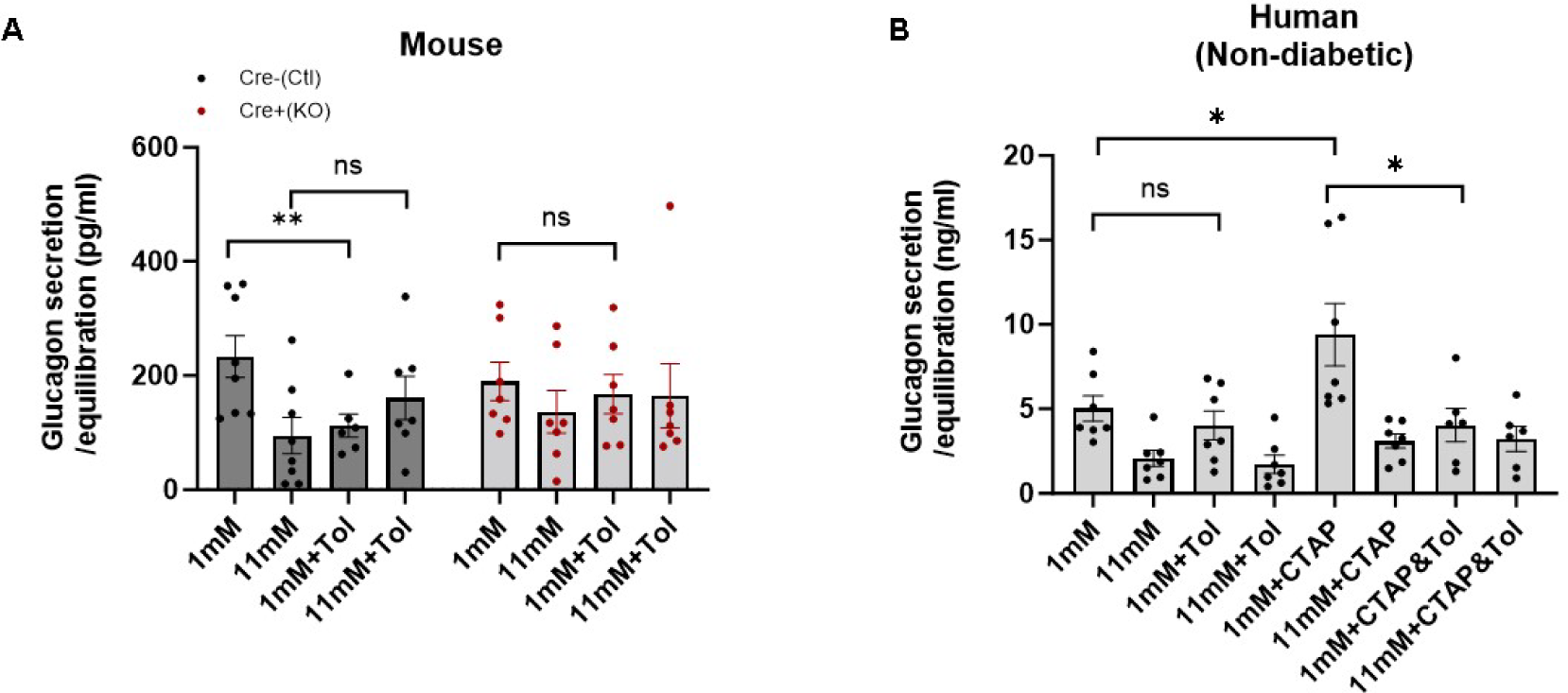
OPRM1 regulates glucagon secretion *in vitro* by a post-KATP mechanism. (A) Tolbutamide (an ATP-sensitive K channel inhibitor) treatment of islets from control and α-cell OPRM1 KO mice. (B) Human islets glucagon secretion assay treated 1 or 11mM glucose with or without 1µM CTAP, 300µM tolbutamide or both (n = 8 experiments from 4 donors). Data represented as means ± SEM. *, *P* < 0.05; **, *P* < 0.01; ***, *P* < 0.001; ****, *P* < 0.0001.

In human islets, the link between K_ATP_ channels and glucagon secretion has been more difficult to parse, perhaps due to the additional recruitment of other ion channels that regulate membrane voltage (e.g., Ca^2+^ channels). Tolbutamide alone has no direct effect on glucagon secretion from human islets. However, K_ATP_ channel function does appear to be necessary for MOPR antagonism-mediated increases in glucagon secretion in low glucose concentrations (Figure 5B). These results could indicate that when K_ATP_ channels are normally expressed, their acute disruption in the context of a disinhibited state (i.e., during MOPR antagonism) reveals an ongoing interaction between K_ATP_ channels and opioid receptors. Together, the mouse and human data suggest that the precise mechanism for MOPR regulation of glucagon secretion involves multiple signaling mechanisms, with the recruitment of K_ATP_ channels appearing to be an important factor.

## DISCUSSION

Disruption of MOPRs in pancreatic islets via either pharmacological antagonism in human islets, or genetic deletion in murine islets, increases glucagon secretion. In contrast, stimulation of MOPRs does not produce measurable effects on glucagon secretion, and neither disruption nor agonism of MOPRs directly influences insulin secretion (Figure 1). There is strong expression of the MOPR gene *Oprm1* in pancreatic α-cells, especially in human islets. MOPR expression is downregulated in metabolically compromised individuals (e.g., T2D), and deletion of the gene in mice predisposes them to a prediabetic phenotype. Finally, while the effects of MOPRs occur via interactions with canonical ion channels like K_ATP_, this is unlikely to be their sole mechanism of action.

A striking result from our experiments is that antagonism of MOPRs yielded a secretory phenotype. In brain, opioids are typically thought to be an evoked system; that is, baseline of occupancy of opioids is thought to be relatively low and thus antagonism alone often does not yield a phenotype. We show that antagonism of human islets or genetic deletion of MOPRs in α-cells of mice is sufficient to potentiate glucagon secretion (Figure 1, 3). This suggests that a functional degree of constitutive activity is present within islets, and this constitutive activity is likely derived from a local, paracrine source [28–30]. Endogenous MOPRs are activated by beta-endorphin or met-enkephalin [7,31]. Unfortunately, literature regarding endogenous opioid peptide expression patterns have been difficult to parse. In some studies, beta-endorphin was shown to be expressed in δ-cells, but in others it was expressed in α-cells [32,33]. Met-enkephalin has been shown to be expressed in β-cells of guinea pig, but whether that translates to mice or humans is unclear [32]. Further complicating the relationship between endogenous opioid peptides and MOPRs are the effects different peptides have on islet activity. For example, both morphine and beta-endorphin increase glucagon secretion [8]; if MOPRs functionally stimulate glucagon secretion, then why would their antagonism also increase glucagon secretion? The simplest explanation is that MOPRs have different roles depending on the islet subtype on which they are expressed. Growing evidence in brain indicates that endogenous opioids may have more nuanced signaling profiles than previously appreciated [19,34,35] (i.e., synaptic rather than volumetric transmission profiles). If this is true for islets, then the precise paracrine interaction likely underlies the seemingly contradictory results. Our data indicate that disruption of MOPRs principally impacts α-cell signaling by increasing glucagon secretion, which may suggest that MOPRs are uniquely constitutively active on α-cells, but not other islet subtypes. Future work using fluorescently labeled receptors or additional measures of receptor signaling effectors should provide further insights into the role of MOPRs on the different islet cell types.

In addition to the hormonal effects of MOPRs on pancreatic islets, our results indicate a relationship between MOPR function and overall metabolic health. We show that islets from either T2D humans or obese mice had downregulated MOPR mRNA relative to healthy controls (Figure 1). We further show that genetic deletion of MOPRs on α-cells did not exhibit health deficits on a chow diet (Supplemental figure 2), indicating that the effects of chronic hyperglucagonemia are potentially dependent on whether the insulin producing β-cells are likewise being taxed [36]. Consistent with this hypothesis, when overall metabolism is challenged by exposure to high-fat diet, MOPR cKO (conditional knock out) mice show impaired glucose tolerance, increased circulating glucagon after glucose administration, increased insulin content, and increased overall islet size (Figure 4). These features are consistent with long-term upregulation of glucagon secretion and associated metabolic diseases like obesity or diabetes [37]. Though not explored in this study, it would be interesting to examine how the long-term upregulation of glucagon secretion from MOPR cKO mice impacts liver function gluconeogenesis or lipid metabolism. Indeed, some evidence points to changes in hepatic responses to glucagon as a major driver of hyperglucagonemia [38,39]. In either individuals with T2D, or in mice with chronically enhanced glucagon secretion, long-term overstimulation of the liver could result in impaired metabolic phenotypes. Complex relationships between liver, alpha, and beta cells could provide an indirect mechanism for alpha cell MOPRs to increase chronically induce hyperglucagonemia and subsequent insulin secretion, eventually leading to insulin intolerance[40]. Collectively, our data support the idea that over stimulation of glucagon secretion and its related effectors (via loss of MOPR function) can drive the transition between a healthy or disease state.

Finally, we sought to determine whether the changes in glucagon secretion observed in our human and MOPR cKO mice were driven by a K_ATP_-dependent mechanism using the selective antagonist tolbutamide. In islets, K_ATP_ channels regulate secretion by exquisitely controlling the membrane potential through the flow of K^+^. This process converges on the exclusion (high K_ATP_ channel activity) or recruitment (low K_ATP_ channel activity) of extracellular Ca^2+^. We find that MOPR cKO mice are unaffected by tolbutamide, such that glucagon secretion remains high in high glucose conditions. In human islets, tolbutamide reduces MOPR antagonism-enhanced glucagon secretion. The difference between mouse and human models could be related to long term changes in paracrine signaling, for example, changes in insulin receptor sensitivity on α-cells [41], or changes in Zn^2+^-mediated inhibition [42]. The latter has been shown to be important for opening K_ATP_ channels, which may be downregulated in MOPR KO mice. Future studies could evaluate acute versus long-term modulation of K_ATP_ channels via MOPR disruption by assessing how chronic opioid administration and/or precipitated withdrawal influences K_ATP_ signaling in islets.

In conclusion, we show that µ opioid receptors directly regulate glucagon secretion in both mouse and human islets via their actions on α-cells. Chronic disruption of MOPRs is associated with metabolic disease states influences neighboring β-cell function. While the molecular mechanism underlying these opioid effects remains unknown, future studies will assess how opioids impact other metrics of metabolic health, as well as how opioids may act on other islet subpopulations.

## SUPPLEMENTAL FIGURES

**Table S1.** Human Islet Donor Demographics, Related to Figure 1. Donor demographics for OPRM1 mRNA expression in human islets.

| Donor ID | Age (years) | gender | BMI | Type 2 Diabetes(+/-) |
| --- | --- | --- | --- | --- |
| SAMN32641505 | 16 | M | 29.5 | - |
| SAMN28673685 | 37 | M | 28.4 | - |
| SAMN33293779 | 49 | M | 68.5 | + |
| SAMN33103085 | 49 | F | 27.5 | - |
| SAMN36020139 | 46 | F | 27.5 | + |
| SAMN37284997 | 62 | F | 43 | + |
| SAMN36823227 | 41 | F | 38.4 | - |
| SAMN37159756 | 34 | M | 33.4 | - |
| SAMN34130383 | 42 | F | 29.2 | - |
| SAMN37350251 | 43 | F | 29.9 | - |

**Supplemental figure 1.**
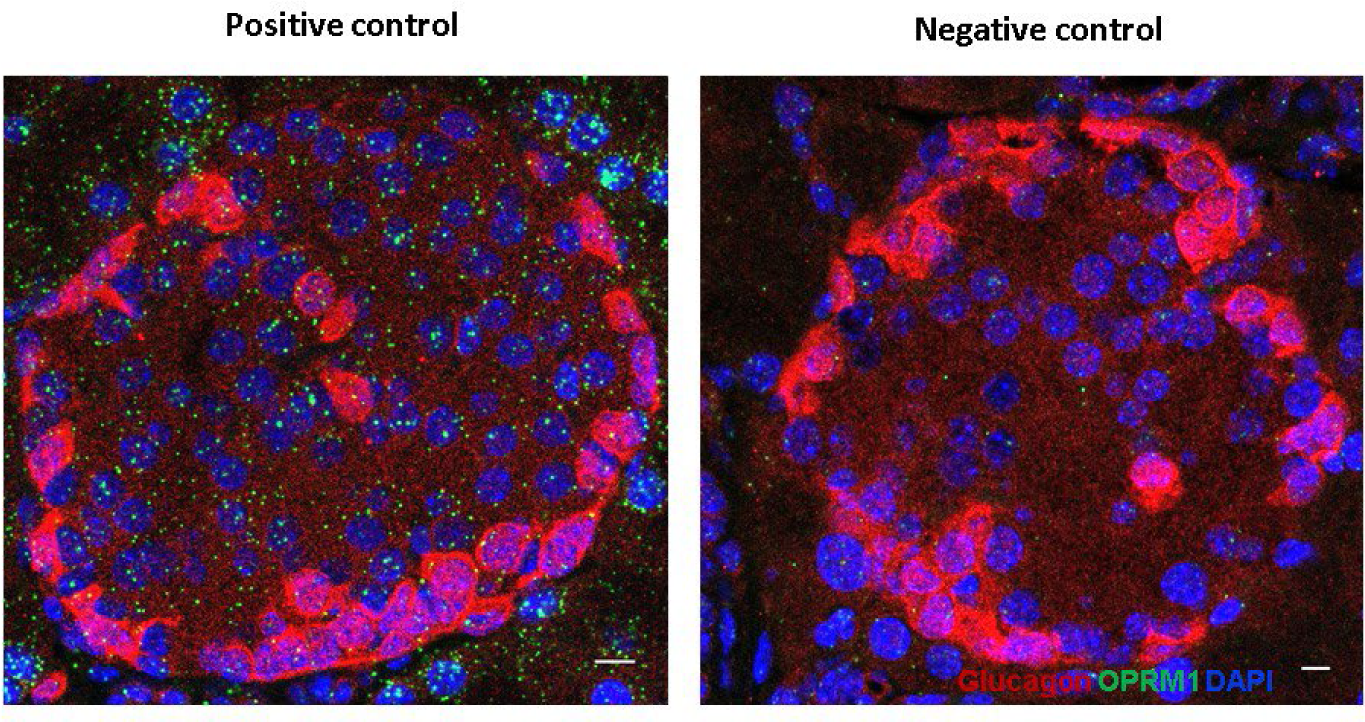
RNAscope controls. RNAscope ISH for positive and negative control probes (green) with immunostaining of glucagon (red) co-detection and nuclei were counterstained with DAPI (blue) in pancreas sections from GCGCre *Oprm1* ^fl/fl^ control mice. Scale bars, 10 μm. Images are representative of three experiments with three mice each.

**Supplemental figure 2.**
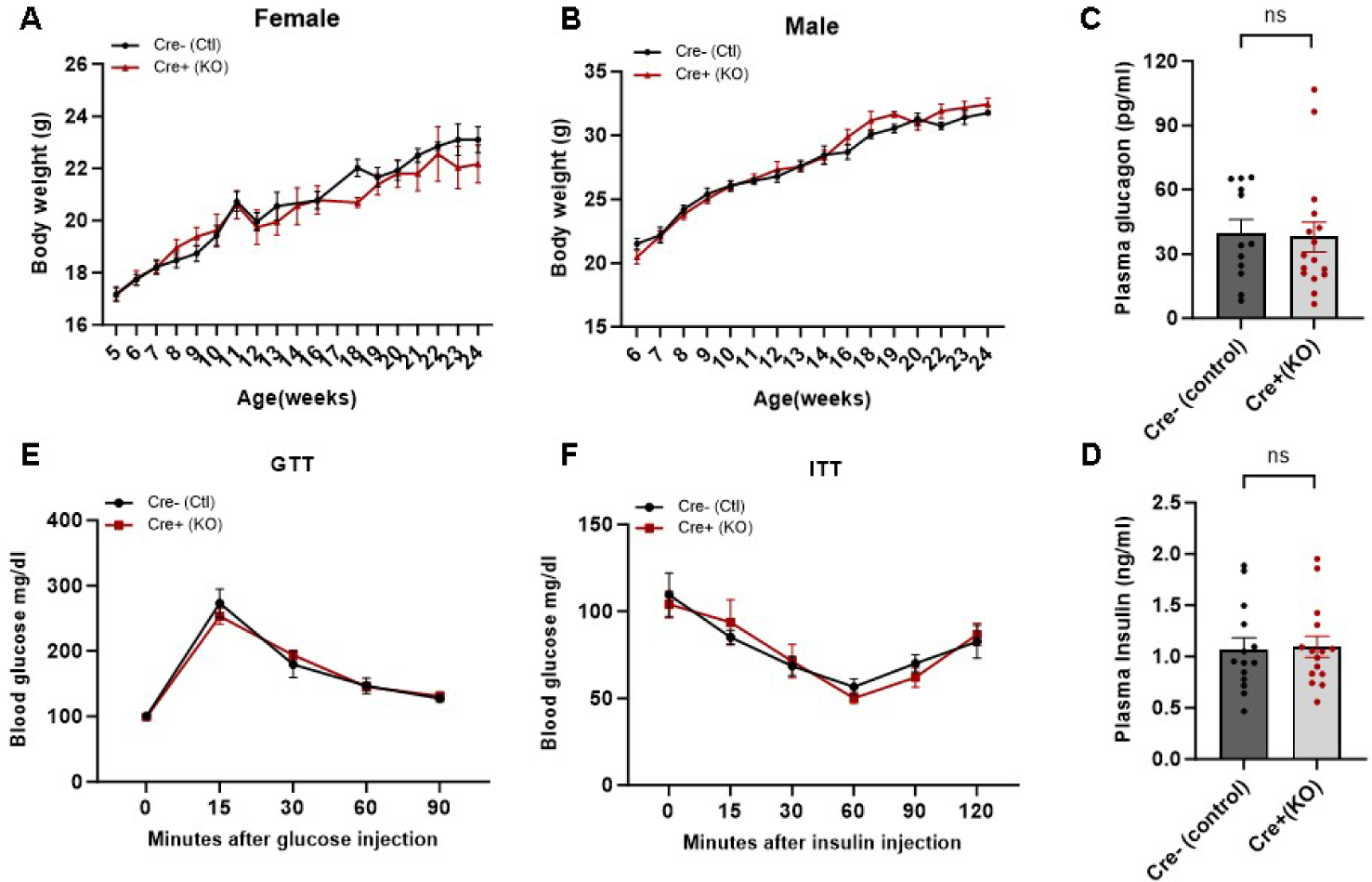
Metabolic phenotypes in chow diet mice. The study had been done in both female and male. Body weight (A) female (B) male, plasma glucagon (C), insulin level (D), GTT (E) and ITT(F) in 10-18 weeks mice are unaffected.

**Supplemental figure 3.**
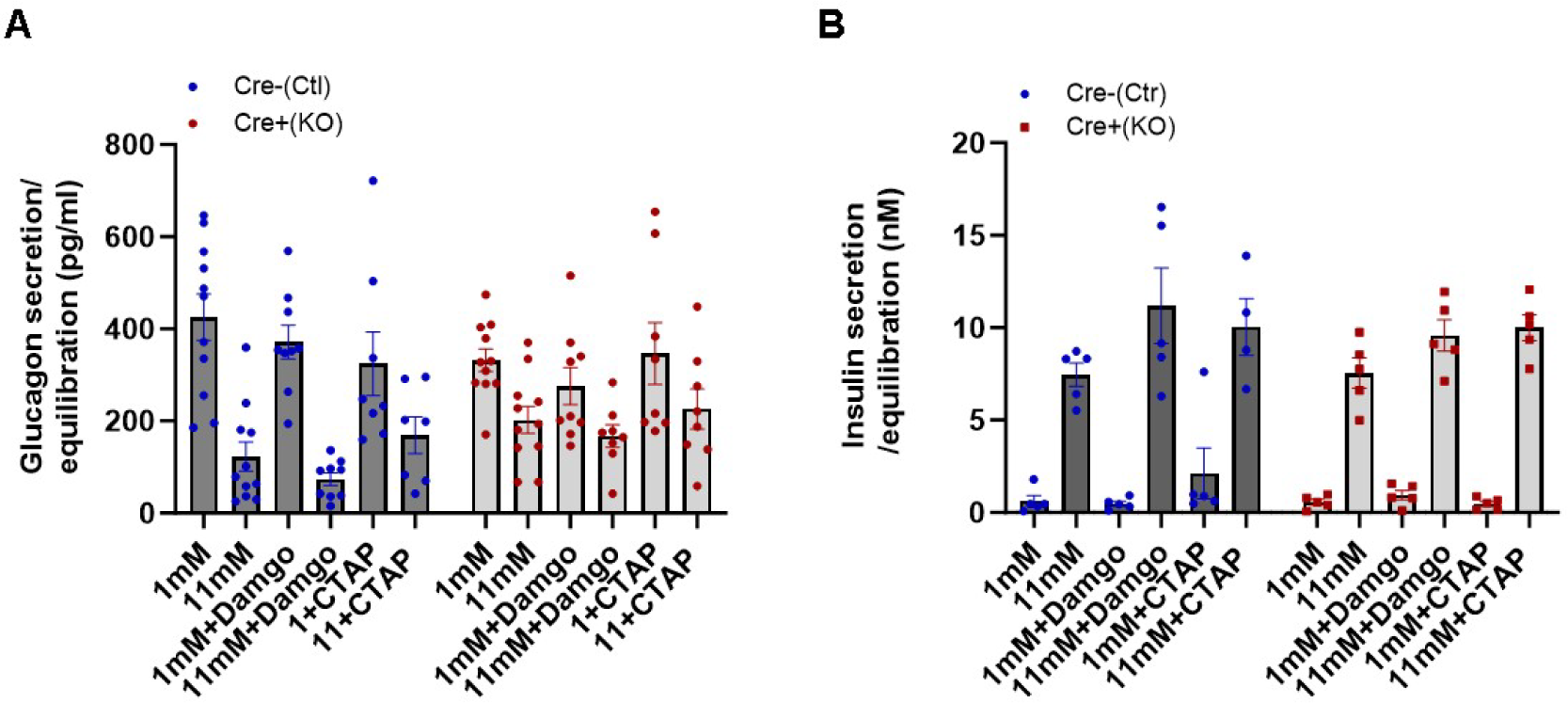
Mouse islet secretion by MOPR agonist DAMGO and antagonist CTAP. Isolated Islet secretion for glucagon (A) and insulin (B) from α cell OPRM1 deficiency (KO) and control (Ctl) mice were treated with receptor agonist DAMGO and antagonist CTAP in low glucose (1mM) and high glucose (11mM).

**Supplemental figure 4.**
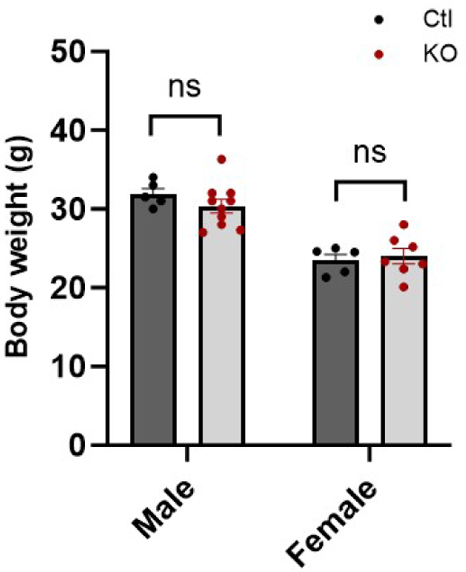
Body weight in high fat diet mice. Body weight of α cell OPRM1 deficiency KO and control mice after 12 weeks on a moderately high fat diet (21.6% calories from fat) in both male and female were measured.

## AUTHOR CONTRIBUTIONS

C.K designed and oversaw all experiments. C.K. D.C.C and J.L planned and performed experiments and interpreted results. D.C.C and C.K wrote the manuscript. All authors reviewed the manuscript.

## DATA AVAILABILITY

Data will be made available on request.

## ACKNOWLEDGMENTS

This work was supported in part by National Institutes of Health Grants R01DK123301 (DWP), R00DA049862 (DCC), R01MH132504 (DCC), and The Leona M. and Harry B. Helmsley Charitable Trust (grants G-2305-06050, G-1912-03558, and G-2001-04215 to DWP). Fluorescence imaging was performed in the Washington University Center for Cellular Imaging (WUCCI) supported by Washington University Diabetes Research Center (P30DK020579).

